# Shared molecular neuropathology across major psychiatric disorders parallels polygenic overlap

**DOI:** 10.1101/040022

**Authors:** Michael J. Gandal, Jillian R. Haney, Neelroop N. Parikshak, Virpi Leppa, Steve Horvath, Geschwind H. Daniel

**Affiliations:** Program in Neurobehavioral Genetics, Semel Institute, David Geffen School of Medicine, University of California, Los Angeles, Los Angeles, CA 90095, USA.; Department of Neurology, Center for Autism Research and Treatment, Semel Institute, David Geffen School of Medicine, University of California Los Angeles, 695 Charles E. Young Drive South, Los Angeles, CA 90095, USA; Department of Human Genetics, David Geffen School of Medicine, University of California, Los Angeles, California, USA; Department of Psychiatry, Semel Institute, David Geffen School of Medicine, University of California Los Angeles, 695 Charles E. Young Drive South, Los Angeles, CA 90095, USA

## Abstract

Recent large-scale studies have identified multiple genetic risk factors for mental illness and indicate a complex, polygenic, and pleiotropic genetic architecture for neuropsychiatric disease. However, little is known about how genetic variants yield brain dysfunction or pathology. We use transcriptomic profiling as an unbiased, quantitative readout of molecular phenotypes across 5 major psychiatric disorders, including autism (ASD), schizophrenia (SCZ), bipolar disorder (BD), depression (MDD), and alcoholism (AAD), compared with carefully matched controls. We identify a clear pattern of shared and distinct gene-expression perturbations across these conditions, identifying neuronal gene co-expression modules downregulated across ASD, SCZ, and BD, and astrocyte related modules most prominently upregulated in ASD and SCZ. Remarkably, the degree of sharing of transcriptional dysregulation was strongly related to polygenic (SNP-based) overlap across disorders, indicating a significant genetic component. These findings provide a systems-level view of the neurobiological architecture of major neuropsychiatric illness and demonstrate pathways of molecular convergence and specificity.

**One Sentence Summary:** Autism, schizophrenia, and bipolar disorder share global gene expression patterns, characterized by astrocyte activation and disrupted synaptic processes.

## Main Text

Decades of neuropathological investigation have failed to identify a consistent neurobiological underpinning of mental illness. Lack of consistent neurobiological findings is often attributed to disease heterogeneity and comorbidity, small sample sizes, limited availability of disease-relevant tissue, and difficulty disentangling the causality of any observed changes. However, the significant heritability of nearly all psychiatric diseases (46% as a class) indicates that genetics will ultimately provide crucial insight into disease mechanisms (*1*). Indeed, recent large genome-wide association (GWA) and exome sequencing studies have robustly identified genetic risk factors, which indicate that the genetic architecture is highly polygenic, with contributions rom many common variants of small effect, as well as rare variants with greater penetrance (*2*-*4*).

Understanding the functional relevance of genetic risk variants, however, remains poorly understood, as few common variants lie within coding regions of the genome and most cannot be functionally linked to a specific gene. This picture is further complicated by significant pleiotropy, variable expressivity and incomplete penetrance (*3*, *5*-*7*). It is now well-established that individual rare variants, such as copy-number variation at 22q11.2 or 16p11.2, can substantially increase risk for multiple disorders including ASD (autism spectrum disorder), schizophrenia (SCZ), and intellectual disability (ID) (*7*). In addition, recent work by the Psychiatric Genomics Consortium (PGC) found significant genetic correlation among SCZ, Bipolar Disorder (BD), and Major Depressive Disorder (MDD), as well as SCZ and ASD using SNP-based co-heritability estimates, indicating shared common genetic variation, as well as shared functional pathways implicated across disorders (*5*, *8*). There is also evidence, however, for some distinct, non-overlapping common genetic risk profiles between SCZ and BD (*9*). As such, a critical remaining question is how polygenic and pleiotropic genetic variants integrate with environmental and epigenetic risk factors in the brain to yield overlapping risk, as well as causing clinically distinct disorders.

We have previously shown that the human brain transcriptome is a highly conserved, robust and hierarchically organized system that can be used as a functional readout of convergent molecular pathology integrating genetic and non-genetic risk factors for heterogeneous neurodevelopmental disorders, such as ASD (*10*-*12*). Here, we leverage this approach to determine whether disease-related transcriptomic signatures are shared across major neuropsychiatric disorders with distinct symptoms and trajectories and whether these patterns reflect common genetic risk factors. This approach is unbiased and relies on carefully correcting for biological and technical covariates, and independent replication, assessing a transcriptomic search space with good coverage, and providing an organizing framework based on gene networks and disease associated genetic variation for the observed changes.

We performed a combined analysis of available gene-expression microarray studies of cerebral cortex across five major neuropsychiatric disorders (*10*, *13*-*20*). A total of 715 brain samples were included from subjects with ASD (n=33), SCZ (n=141), BD (n= 75), MDD (n=68 samples), alcohol abuse disorder (AAD, n=17), and matched non-psychiatric controls (n=381) (see **Methods**). Inflammatory bowel disease (IBD, n=67) was included as a non-neural comparison (*21*). These disorders are highly prevalent (ranging from 0.6% to 17%) and disabling, collectively accounting for 43% of global disease burden associated with neurological, psychiatric, and substance abuse disorders (*22*). Individual datasets underwent identical, strict quality control and normalization procedures (Figure 1; **Methods**), including re-balancing to remove any confound between diagnosis and biological (e.g., age, sex) or technical (e.g., postmortem interval, pH, RIN, batch, 3’ bias) covariates. Probes were re-annotated using a standardized nomenclature (Ensembl v75; **Methods**), and experimental batch effects were corrected both within and between studies (*23*).

**Figure 1.**
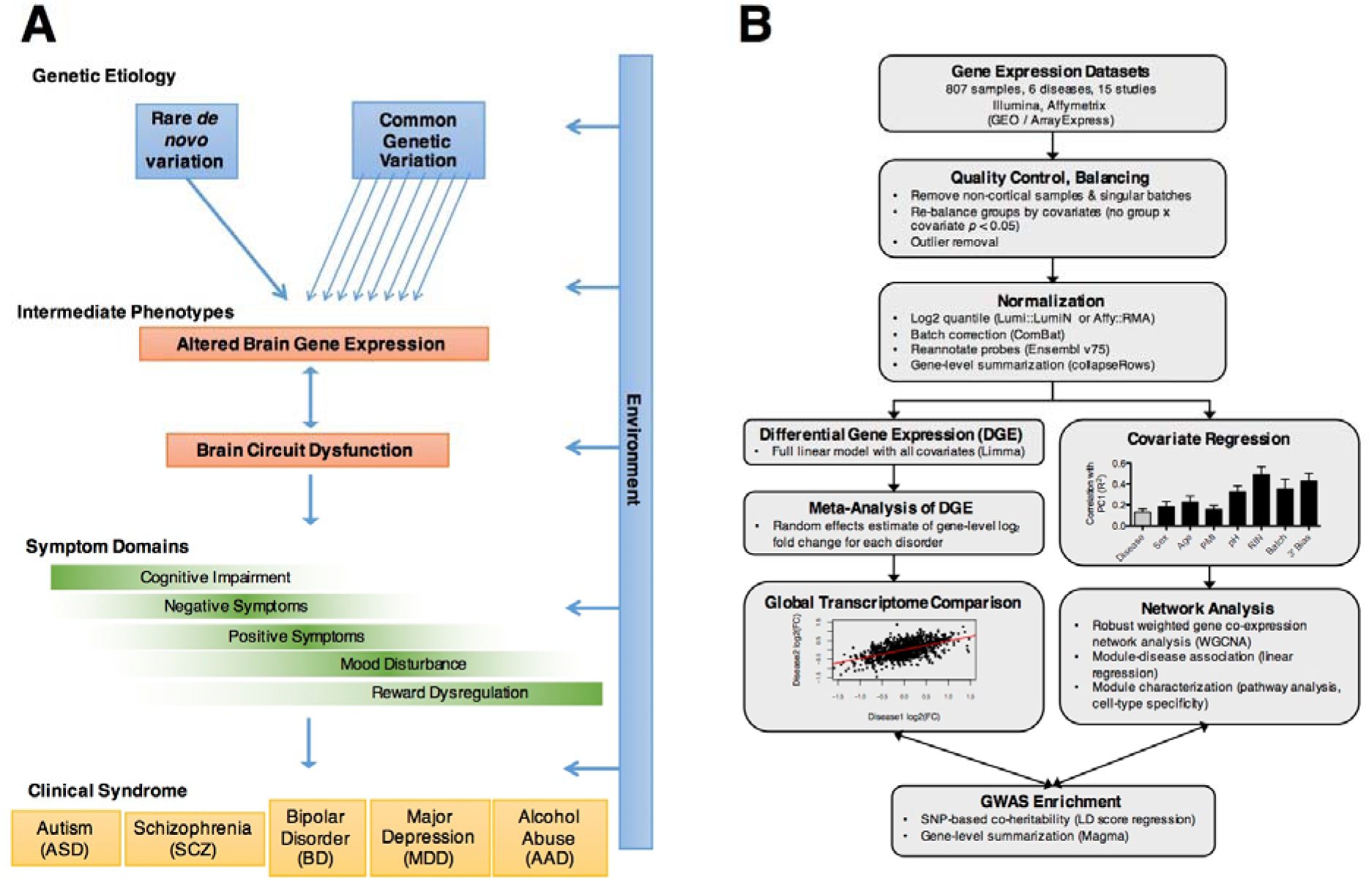
**A)** Model of psychiatric disease pathogenesis. **B)** Cross disorder transcriptome analysis pipeline (see **Methods**). Cortical gene expression datasets were compiled from cases of ASD (n=33 samples), schizophrenia (n=141), bipolar disorder (n=75), depression (n=65), alcoholism (n=17), and matched non-psychiatric controls (n=381). Datasets underwent identical, strict quality control and normalization procedures, including outlier removal and rebalancing to remove any group x covariate confound. Probes were mapped to a standardized nomenclature (Ensembl v75; hg19) and collapsed to the gene level. Differential gene expression was calculated using all available biological (disorder, sex, age) and technical (PMI, pH, RIN, batch, 3’ bias) covariates. Network analysis (WGCNA) was performed on covariate-regressed data. Gene coexpression modules were assessed for disease association, cell-type specificity, functional pathways, and enrichment for GWAS signals.

Differential gene expression (DGE) signatures were computed as log_2_ fold-change (log_2-_FC) values for each case/control comparison using a linear regression framework in *limma*, accounting for all available covariates (*24*). A random-effects meta-analytic approach (restricted maximum-likelihood estimates) was taken to compute disease-specific DGE signatures for each disease (*25*). Spearman’s correlation of disease DGE signatures revealed a significant, global transcriptome overlap among ASD, schizophrenia, and bipolar disorder as well as among schizophrenia, bipolar disorder, and depression (Figure 2A; all rho ≥ 0.2, *P* < 10^-15^, Bonferroni corrected). The slopes of linear regressions between ASD, BD, and MDD vs SCZ were 1.51, 0.75, and 0.20, indicating a gradient of transcriptomic severity with ASD > SCZ > BD > MDD (**Figure S1**). Interestingly, ASD and IBD shared an overlap with rho 0.15 (*P* < 10^-15^), likely due to common inflammatory changes (*26;* see also below). The lack of (or negative) overlap between chronic alcoholism and other disorders suggests that similarities are less likely due to comorbid substance abuse, poor overall general health, or general brain-related post-mortem artifacts.

**Fig. 2.**
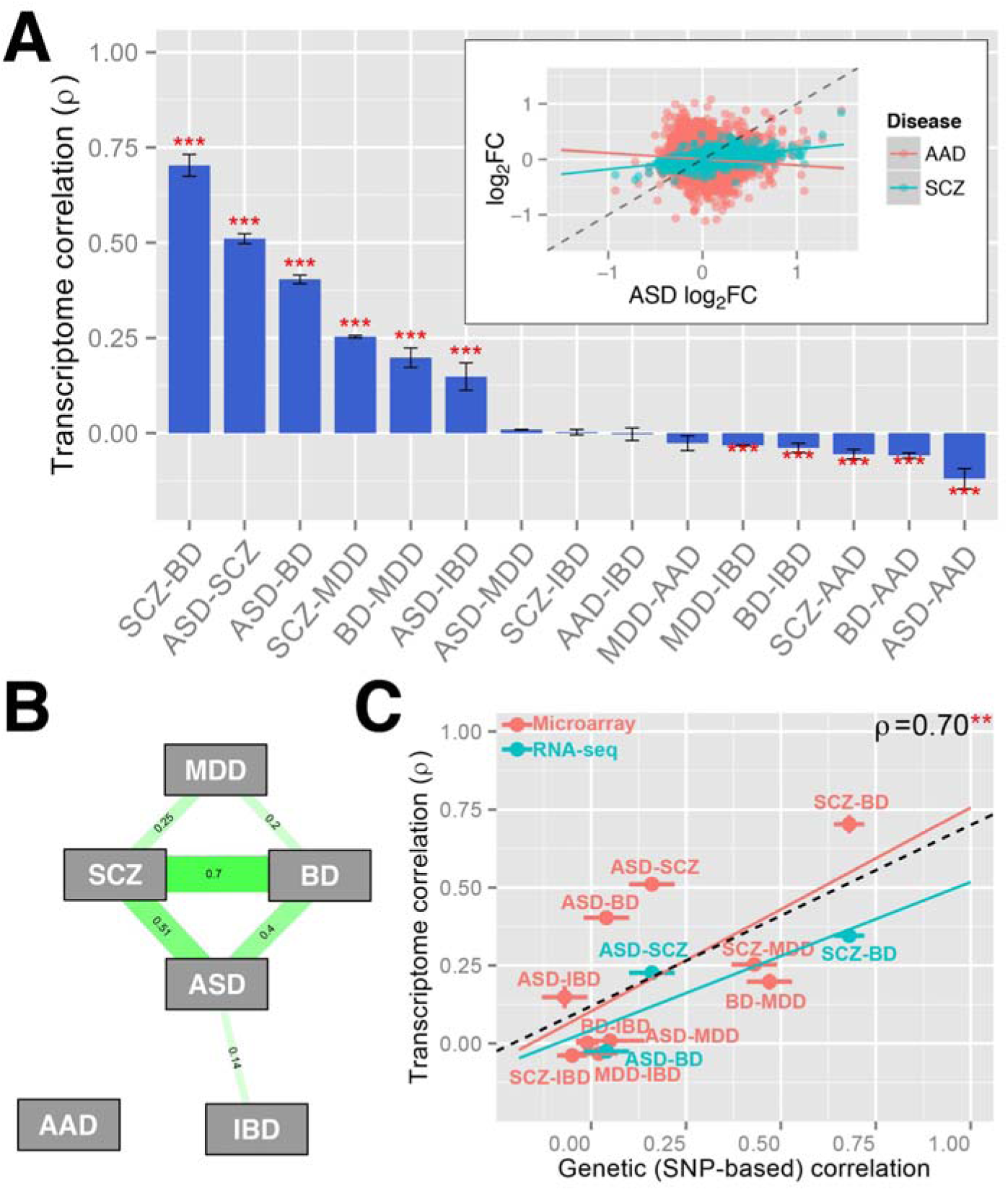
Cortical gene expression patterns overlap across distinct psychiatric disorders and share common genetic risk. **A)** Rank order of transcriptome similarity for all disease pairs, as measured by spearman’s correlation of differential expression log_2_ fold-change values. Significant overlap is observed between ASD, SCZ, and BD as well as between SCZ, BD, and MDD (all *ρ* ≥ 0.2, *P* < 10^-15^, Bonferroni corrected). Inset, differential expression values are plotted for significant (ASD-SCZ) and non-significant (ASD-AAD) disease pairs. Chronic alcoholism shows no positive overlap with any other disorder. **B)** Overlapping gene expression patterns across disease are strongly correlated with shared common genetic variation, as measured by SNP-based co-heritability (*5*). A similar pattern is observed for microarray-based (discovery) and RNA-seq (replication) datasets. ** *P* < 0.01, *** *P* < 10^-15^ (Bonferroni corrected).

Disease-specific DGE signatures included in **Table S1** provide benchmarks for determining the relevance of model organisms, *in vitro* systems (e.g., iPSC), or drug effects for matching *in vivo* human disease relevant transcriptomes (*27*, *28*). We note that the most concordantly downregulated genes (**Figure S2**) include the cortical interneuron markers parvalbumin (*PVALB) and somatostatin (SST)*, corticotrophin releasing hormone (*CRH)*, inducible nerve growth factor (*VGF*), the presynaptic vesicle membrane protein synaptogyrin (*SYNGR3*), and the neurogenic transcription factor *NEUROD6*. Genes most upregulated across disease included both dimers of calprotectin (*S100A8* and *S100A9*). To investigate the potential contribution of psychiatric medication, disease-specific transcriptome signatures were compared with gene expression changes in non-human primates treated with acute or chronic antipsychotic medications that are commonly used in schizophrenia, showing significant negative overlap (**Figure S3**), indicating that the observed patterns were not caused by these medications.

To validate that these transcriptomic phenotypes were reproducible, we used independent RNAseq datasets for replication, which although smaller than the microarray datasets, were available for 3 out of the 5 disorders. The first consisted of 100 bp paired end ribosomal depleted reads from prefrontal cortex samples of schizophrenia (n=275), bipolar (n=47), and matched controls (n=262) as part of the CommonMind Consortium (commonmind.org). A second dataset consisted of 100bp single end reads from frontal and occipital tissue from subjects with ASD (n=32) and matched controls (n=40) (*29*). Reads were aligned and mapped according to a standardized workflow (**Methods**) and underwent conditional quantile normalization to correct for biases introduced due to read depth, gene-length, and GC content. Differential expression was assessed using a multiple linear regression framework using all available covariates including multiple RNAseq alignment and mapping statistics to control for technical variation. Comparison of DGE signatures across disorders revealed very similar results to microarray analyses, with strong overlap among ASD, SCZ, and BD (see Figure 2B).

### Co-Expression Networks Refine Axes of Psychiatric Pathophysiology

To characterize the biological pathways overlapping across disorders, we performed weighted gene co-expression network analysis (WGCNA) on the entire set of (covariate-regressed and batch-corrected; **Methods**) cases and controls. We used robust WGCNA (*30*-*32*) to avoid the influence of potential outlier samples or studies. Modules were identified using unsupervised hierarchical clustering based on topological overlap of signed networks and annotated based on enrichment of gene ontology pathways, as well as markers of specific cell-types, development time-points, and regional identity within the central nervous system (Figure 3, **S4**). Several disease-associated modules were highly enriched for markers of specific CNS cell types including neurons, astrocytes, microglia and endothelial cells. Cross-disorder (CD) modules were assigned colors and numbers as identifiers. The gene expression levels of a module are represented by the module eigengene, which is defined as the first principle component of the standardized expression values (11). Differential eigengene expression was quantified in two complementary ways – linear regression of module eigengene with disease, and overlap between up or down-regulated gene lists for each disorder (FDR < 0.05) and each module.

**Fig. 3.**
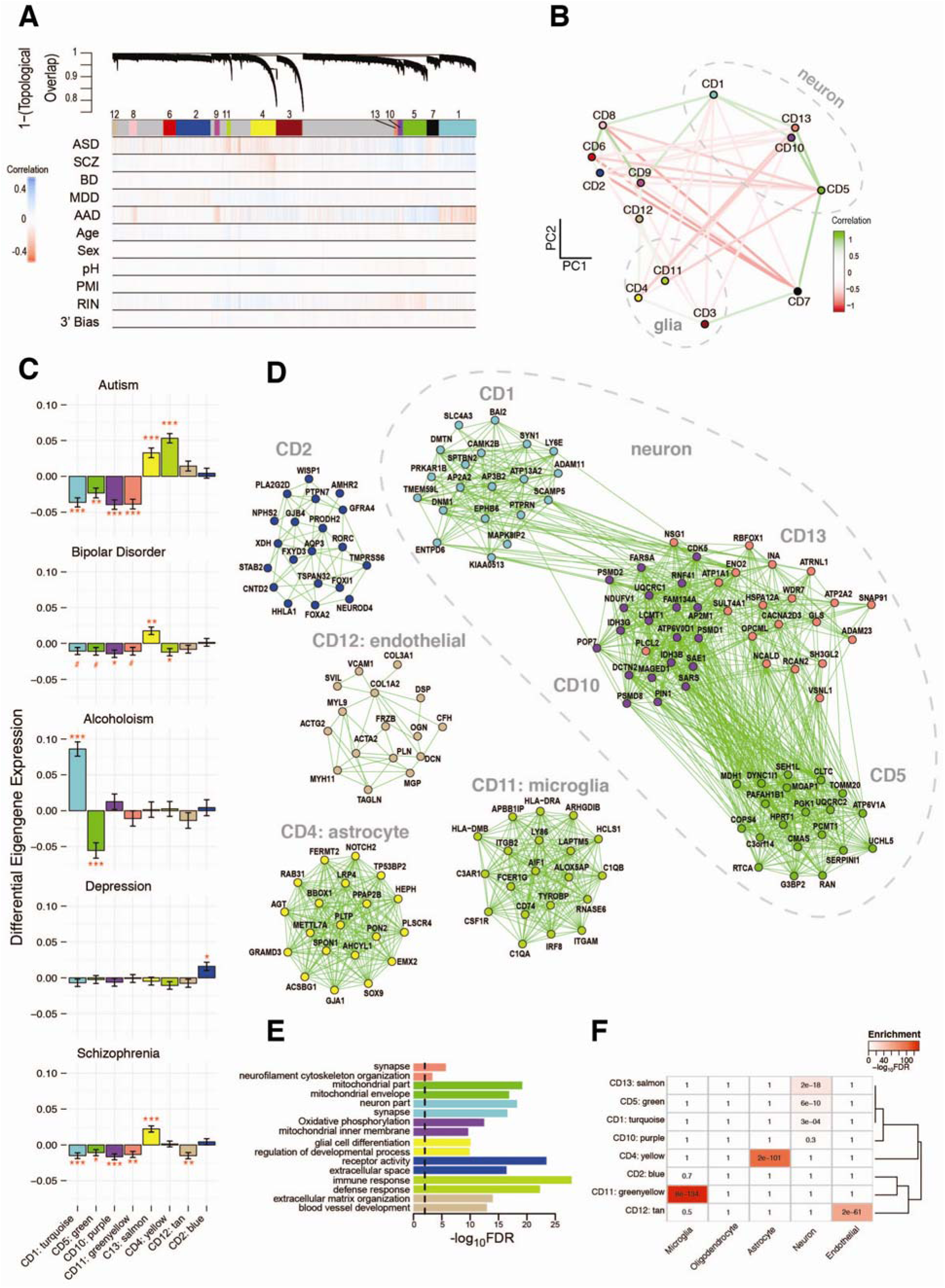
Network analysis identifies modules of coexpressed genes dysregulated across disease in distinct combinations. **A)** Network dendrogram is based on co-expression topological overlap of genes across disorders. Color bars show correlation of gene expression with disease status, biological, and technical covariates. **B)** Multidimensional scaling plot demonstrates relationship between modules and clustering by cell-type relationship. **C)** Module eigengene differential expression is perturbed across disease states. Plots show beta values from linear regression beta values of module eigengene with disease status. Upregulated modules reflect G-protein coupled receptors in MDD, microglia in ASD, and astrocytes in ASD, SCZ, and BD. Downregulated modules reflect neuronal/synaptic and mitochondrial processes across multiple disorders. FDR-corrected p-values are highlighted (#*P*<0.1, \**P*<0.05, \*\**P*<0.01, \*\*\**P*<0.001). **D)** The top twenty hub genes are plotted for modules most disrupted in disease. See **Table S1** for a complete list of genes’ module membership. Edges are weighted by the strength of correlation between genes. Modules are characterized by (**E)** Gene Ontology enrichment (top two pathways shown for each module) and (**F)** cell-type specificity, based on RNA-seq of purified cell populations from healthy human brain samples (*67*).

Module CD4 (yellow) was enriched for astrocyte markers and significantly upregulated in ASD (FDR-corrected *P*=9.5 × 10^-6^; **Methods**), BD (FDR-corrected *P*=0.0029), and SCZ (FDR corrected *P*=6.5 ×10^-7^; Figure 3C), consistent with previous reports of astrocyte activation in schizophrenia (*33*) and ASD (*34*). Pathways enriched among the genes contained within this module included glial cell differentiation as well as organic acid, fatty-acid, and lipid metabolism. The most highly connected genes within this module, or hub genes, included the gap junction protein *GJA1* (e.g., connexin-43), *PLTP*, a critical component of the blood-brain barrier, and *SPON1*, encoding an extracellular matrix protein, which was recently identified as a genome-wide significant regulator of brain structure and connectivity (*35*).

In contrast, module CD11 (greenyellow) was enriched for microglial markers and was significantly upregulated only in ASD (FDR-corrected *P*=7.5 × 10^-14^). Hubs include major components of the complement system (*C1QA, C1QB*), canonical microglial markers (*HLA-DRA* and *AIF-1*), and *TYROBP*, a key regulator of late-onset Alzheimer’s disease neuropathology (*36*,*37*). Results agree with convergent evidence for neuroimmune/inflammatory dysfunction in ASD (*38*), as well as emerging understanding of the importance of microglia in the regulation of synaptic development and function (*39*). Of note, microglial activation has been previously reported in SCZ and, to lesser extent, in BD. However, these studies have been small and often fail to account for important covariates, leading to mixed results (*40*). Furthermore, recent work has demonstrated that PET imaging ligands used as *in vivo* markers for activated microglia also cross-react with activated astrocytes (*41*). These results suggest that microglial activation is not present to the same magnitude in SCZ and BD as is seen in ASD, or at the level of astrocyte upregulation that is observed across all three disorders.

Module CD2 (blue) was upregulated only in MDD (FDR-corrected *P*=0.011) and did not show strong cell-type enrichment. Pathways enriched in this module included G-protein coupled receptors, cytokine-cytokine interactions, and glucocorticoid metabolic process. Hub genes include *LHX3*, a transcription factor critical for pituitary development and hormone signaling (*42*), and *FOXA2*, an insulin target that regulates levels of orexin and melanin-concentrating hormone (MCH) and is a critical regulator of serotoninergic and dopaminergic neurogenesis (*43*). Notable genes in this module also included the neuropeptide oxytocin (*OXT*), both melatonin receptors (*MTNR1A, MTNR1B*), thyrotropin-releasing hormone and its receptor (*TRH*, *TRHR*), the ACTH receptor (*MC2R*), the orexin receptor *HCRTR1*, several serotonin receptors (*5HTR-1B*, *5HTR-1D*, *5HTR-3A*), and dopamine receptors (*DRD2*, *DRD3*). Together, these genes indicate a link between inflammation, dysregulation of the hypothalamic-pituitary axis, and monoaminergic neurmodulation, supporting current models of MDD pathophysiology (*44*). Furthermore, these pathways provide specific molecular connections to symptomatic disturbances in sleep-wake regulation, appetite, and energy balance that are observed in patients with depression.

Two modules containing neuronal markers (CD1, turquoise; CD13, salmon) were significantly down-regulated in ASD (CD1: FDR-corrected *P*=6.5×10^-8^; CD13: FDR-corrected corrected 2.6×10^-7^),and SCZ (CD1: FDR-corrected *P*=5.2×10^-4^; CD13: FDR-corrected *P*=7.2×10^-3^).These modules significantly overlapped with downregulated genes in BD (CD1: OR 7.4, FDR-corrected *P*=5×10^-11^; CD13: OR 5.5, FDR-corrected *P*=0.043), although the eigengene-BD correlation was not significant (CD1: FDR-corrected *P*=0.074; CD13: FDR-corrected *P*=0.106). Although the same genes were dysregulated across the three disorders, ASD showed the most extreme changes within these shared modules, suggesting a more severe molecular phenotype, and perhaps related to its earlier clinical onset. CD13 was enriched for gene ontology pathways including synapse, neuron, and calcium mediated signalling. Notable genes in this shared module included the RNA splicing regulator *RBFOX,* as well as the calcium binding protein parvalbumin (*PVALB)* and several GABA_A_-receptor subunits. Pathways associated with CD1 included synaptic transmission, voltage gated cation channel, and cholinergic synapse. CD1 module hub genes included *CAMK2B*, an important regulator of synaptic plasticity, and *DNM1*, a GTPase that plays an important role in synaptic vesicle recycling and is linked to epileptic encephalopathy. CD1 also included multiple GABA_B_ receptor subunits, as well as genes involved in syndromic forms of ASD (*mTOR*, *TSC1*, *TSC2*, *MECP2*).

Two remaining modules, CD5 (green) and CD10 (purple), also showed neuronal enrichment with functional annotation most driven by mitochondrial processes. CD5 was downregulated in AAD (FDR-corrected *P*=1.1×10^-6^), ASD (FDR-corrected *P*=0.0023), and SCZ (FDR-corrected *P*=0.029), and included the inhibitory interneuron marker calbindin, both of the rate-limiting enzymes required for GABA synthesis (*GAD1*, *GAD2*), as well as several GABA_A_, glycine, and glutamate receptors. CD10 was downregulated in ASD (FDR-corrected *P*=4.7×10^-8^), SCZ (FDR-corrected *P*=3.5×10^-4^), and BD (FDR-corrected *P*=0.023), and characterized by pathways including oxidative phosphorylation and respiratory electron transport chain. Hub genes include *ATP6V0D*, a proton pump important for CNS patterning, as well as the peptide hormone and interneuron marker *CCK* as well as its receptor *CCKBR*. These results provide further evidence for the link between energetic balance, synaptic transmission, and psychiatric disease (*45*).

The transcriptome represents a brain phenotype that may either reflect causal factors, or the consequences of the disorder. To assess the relationship of transcriptional alterations to causal genetic factors, we next asked whether overlapping transcriptome patterns across disorders are related to shared genetic aetiologies, as measured by SNP-based polygenic correlation from a recent cross disorder analysis of GWAS studies performed by the PGC (*5*). Remarkably, common genetic correlation between psychiatric disorders was significantly associated with transcriptome similarity across the same disease pairs (Figure 2B; Spearman’s *ρ*=0.70, 95% confidence interval [0.25–0.90], *P*=0.007), indicating that the effect of common variants associated with disease is also captured in cortical gene-expression patterns

To further determine which shared and distinct cross-disorder biological processes were related to genetic risk factors, we investigated enrichment of both common and rare disease-associated variants within dysregulated modules. Recent estimates suggest that common genetic variation may account for up to 50% of the heritability associated with these disorders (*47*, *48*). We used MAGMA, a multiple regression framework for gene-based analysis that explicitly accounts for linkage disequilibrium between SNPs (*49*), to generate aggregate gene-level significance values from the most recent GWAS studies of schizophrenia, bipolar, ASD, alcoholism, depression, and inflammatory bowel disease, the latter as a non-neural comparison (*2*, *50*-*54*). We quantified the relationship between a gene’s module membership (kME), a relative measure of centrality or hubness, and GWAS-derived gene significance for each module using Spearman’s correlation. The most significant signal comes from the latest schizophrenia GWAS, the most highly powered of such studies, which demonstrates significant enrichment of risk genes within all four down-regulated neuronal modules (Figure 4A; CD1, CD5, CD10, CD13 all FDR-corrected P-values < 10^-14^). In aggregate, these modules account for 3.3% of the variance in GWAS signal in schizophrenia, which is small, but of biological significance. Enrichment within three of these neuronal modules was also observed for the most recent PGC bipolar GWAS (CD1: FDR-corrected *P*=6.8×10^-6^; CD10: FDR-corrected *P*=0.043; CD13: FDR-corrected *P*=0.027; Figure 4A). No significant enrichment was found for ASD or alcoholism GWAS results, which may reflect lower power due to their relatively small sample sizes. Finally, none of the microglial or astrocyte specific modules showed polygenic GWAS enrichment, suggesting a non-genetic etiology underlying these processes, as previously suggested in ASD (*10*).

**Fig. 4.**
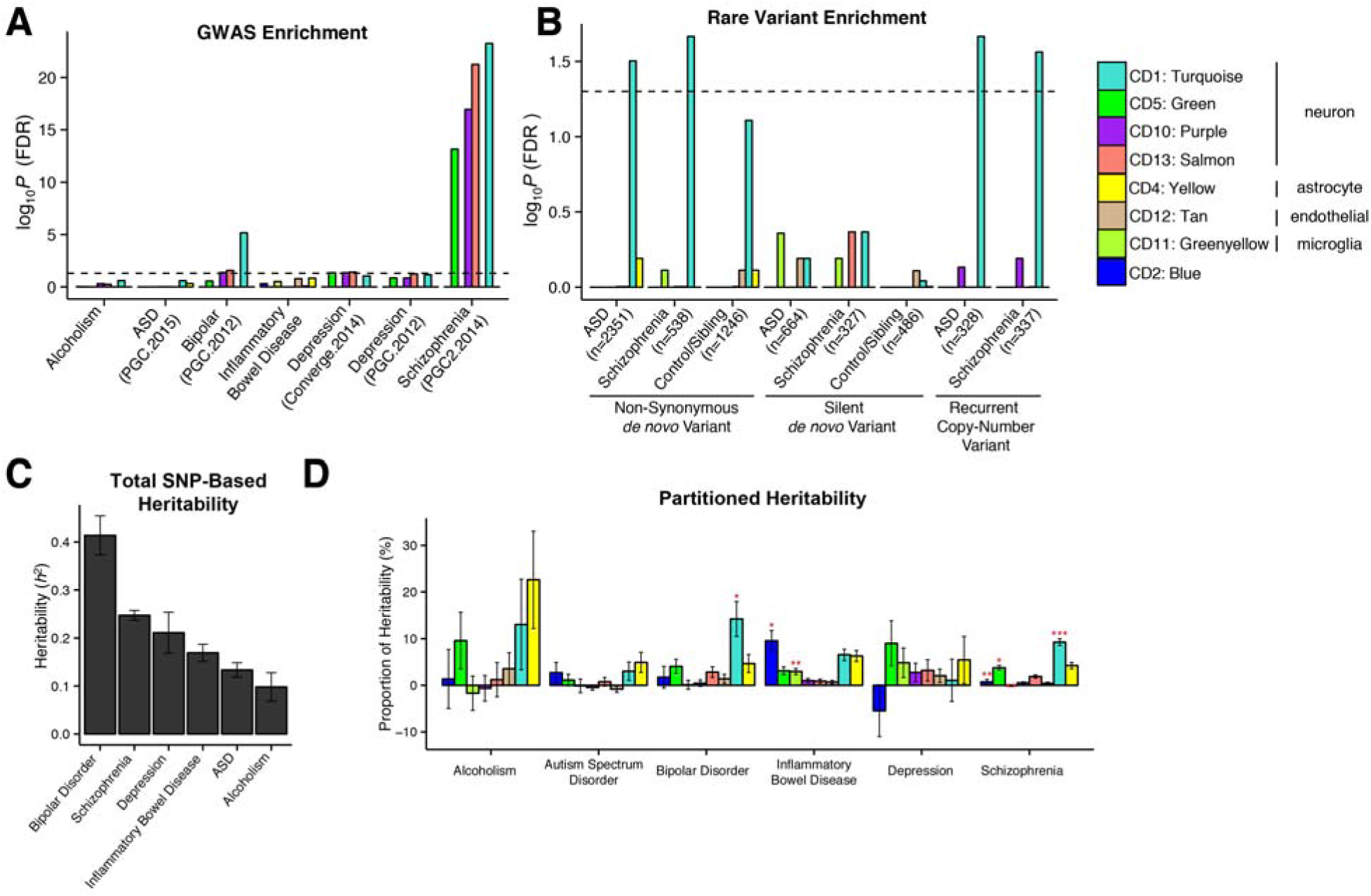
Neuronal modules downregulated in ASD, SCZ, and BD are enriched for common and rare genetic risk factors. **A)** Significant enrichment is observed for SCZ‐ and BD-associated common variants from GWAS among neuron/synapse & mitochondrial modules (P-values are FDR-corrected). Gene-level significance values were calculated from GWAS summary statistics (*2*, *50*-*54*) using magma (*49*). For each module, spearman’s correlations were used to assess the relationship between gene module membership (kME) and GWAS significance (-log_10_P-value). **B)** The CD1 (turquoise) neuronal module shows significant enrichment for non-synonymous *de novo* variants identified in recent whole exome sequencing studies (*3*, *58*-*64*). No enrichment is seen in unaffected controls or siblings, nor was enrichment seen in genes affected by silent mutations. CD1 is also enriched for genes affected by recurrent copy-number variation in ASD and SCZ (see **Table S4**). The number (n) of genes affected by different classes of variation are shown in parentheses. Significance was calculated using logistic regression, accounting for gene-length as a covariate. P-values are FDR corrected. **C)** Total SNP-based heritability (liability-scale) calculated from GWA studies in (**A**). **D)** Proportion of heritability for each disorder that can be attributed to each module. Significance (*p<0.05,**p<0.01, ***p<0.001, FDR-corrected) is based on enrichment statistics comparing the proportion of SNP heritability within the module divided by the proportion of total SNPs represented. The CD1 (turquoise) module shows significant enrichment in SCZ and BD, accounting for ∼10% of heritability within each dataset despite only containing ∼3% of the SNPs.

There has been some suggestion that rare, *de novo* genetic variants associated with psychiatric disease disrupt different molecular pathways than common variants. For example, rare CNVs tend to be associated with reduced intelligence measures even in subjects without psychiatric diagnoses (*55*) whereas sub-threshold accumulation of common genetic variants tends to be associated with improved cognitive outcomes (*56*, *57*). We assessed for overlap between modules and genes containing non-synonymous or silent *de novo* mutations compiled from recent exome-sequencing studies of ASD, SCZ, and unaffected controls (Figure 4B) (*3*, *58*-*64*). Remarkably, the CD1 neuronal module was enriched for non-synonymous rare variants identified in ASD (OR 1.58, FDR-corrected *P*=0.03) and SCZ cases (OR 1.74, FDR-corrected *P*=0.011), but not in unaffected siblings or controls. No enrichment was found for silent mutations, which serve as a control comparison. A similar pattern was seen for genes within regions affected by recurrent copy-number variation in ASD (OR 2.42, FDR-corrected *P*=0.025) and SCZ (OR 2.34, FDR-corrected *P*=0.032). These results point to a convergence of common and rare genetic risk factors acting to downregulate a specific group of co-regulated genes involved in synaptic transmission across multiple neurodevelopmental psychiatric disorders.

To complement this analysis, and further extend it to common variation, we used LD score regression (*65*) to partition disease heritability (**Methods**; Figure 4D) into the specific contribution from SNPs located within genes from each module. The neuronal module CD1 again showed significant enrichment within both SCZ (enrichment 2.48 fold, FDR-corrected *P*<10^-13^) and BP (enrichment 3.9 fold, FDR-corrected *P*<0.004) GWAS, accounting for ∼10% of SNP-based heritability within each dataset despite only containing 3% of the SNPs. This illustrates how biologically meaningful co-expression modules can be used to understand how a complex pattern of many common variants, each of low effect size, can be integrated with network analysis to implicate specific biological roles for common variant risk across neuropsychiatric disorders.

In conclusion, these data provide a systematic, quantitative, and genome-wide characterization of the molecular alterations in the cerebral cortex of five major neuropsychiatric disorders. We demonstrate how such analysis provides a framework for understanding the common, downstream molecular signalling pathways underlying major neuropsychiatric illness and for interpreting gene variants implicated in disease risk. We observe a gradient of synaptic gene down-regulation, with ASD > SZ > BD. BD and SCZ appear most similar in terms of synaptic dysfunction and astroglial activation and are most differentiated by subtle downregulation in microglial and endothelial modules. ASD shows the most pronounced upregulation of a microglia signature, which is minimal in SCZ or BD. Based on these data, we hypothesize that a more severe synaptic phenotype, as well as the presence of microglial activation, is responsible for the earlier onset of symptoms in ASD, compared with the other disorders, consistent with an emerging understanding of the critical non-inflammatory role for microglia in regulation of synaptic connectivity during neurodevelopment (*39*, *66*). MDD shows neither the synaptic nor astroglial pathology observed in SCZ, BD. In contrast, in MDD, a striking dysregulation of HPA-axis and hormonal signalling not seen in the other disorders is observed. These results provide the first systematic, transcriptomic framework for understanding the pathophysiology of neuropsychiatric disease, placing disorder-related alterations in gene expression in the context of shared and distinct genetic effects.

## Acknowledgments

The work is funded by the US National Institute of Mental Health (P50-MG106438, DHG; R01-MH094714, DHG), the Simons Foundation for Autism Research (SFARI 206733, DHG), and the Stephen R. Mallory schizophrenia research award at UCLA (MJG). The authors thank Hyejung Won, Jason Stein, Damon Poliodakis, Chris Hartl, Jonathan Flint, Bogdan Pasaniuc, Roel Ophoff and members of the Geschwind laboratory for critical reading of this manuscript. The authors thank Bogdan Pasaniuc for assistance with module heritability analyses. Replication RNAseq data were generated as part of the CommonMind Consortium supported by funding from Takeda Pharmaceuticals Company Limited, F. Hoffman-La Roche Ltd and NIH grants R01MH085542, R01MH093725, P50MH066392, P50MH080405, R01MH097276, RO1-MH-075916, P50M096891, P50MH084053S1, R37MH057881 and R37MH057881S1, HHSN271201300031C, AG02219, AG05138 and MH06692. Brain tissue for the study was obtained from the following brain bank collections: the Mount Sinai NIH Brain and Tissue Repository, the University of Pennsylvania Alzheimer’s Disease Core Center, the University of Pittsburgh NeuroBioBank and Brain and Tissue Repositories and the NIMH Human Brain Collection Core. CMC Leadership: Pamela Sklar, Joseph Buxbaum (Icahn School of Medicine at Mount Sinai), Bernie Devlin, David Lewis (University of Pittsburgh), Raquel Gur, Chang-Gyu Hahn (University of Pennsylvania), Keisuke Hirai, Hiroyoshi Toyoshiba (Takeda Pharmaceuticals Company Limited), Enrico Domenici, Laurent Essioux (F. Hoffman-La Roche Ltd), Lara Mangravite, Mette Peters (Sage Bionetworks), Thomas Lehner, Barbara Lipska (NIMH).

## Supplementary Materials

### Materials and Methods

Figures S1-S4

• Figure S1 – Gradient of Transcriptomic Severity

• Figure S2 – Top differentially expressed genes across disorders

• Figure S3 – Antipsychotic associated gene expression changes in non-human primates.

• Figure S4 – Brain regional identity for co-expression modules

Tables S1-S5

• Table S1 – Disease-specific differential gene expression signatures and kME table

• Table S2 – Module Gene Ontology enrichment

• Table S3 – Module eigengene associations

• Table S4 – Compilation of de Novo mutations and CNVs

• Table S5 – Partitioned heritability

Supplemental References

